# The type of carbon source not the growth rate it supports can determine diauxie

**DOI:** 10.1101/2023.10.18.562896

**Authors:** Yu Huo, Weronika Danecka, Iseabail Farquhar, Kim Mailliet, Tessa Moses, Edward W. J. Wallace, Peter S. Swain

**Affiliations:** Centre for Engineering Biology, University of Edinburgh; School of Biological Sciences, University of Edinburgh; EdinOmics, RRID:SCR_021838, Centre for Engineering Biology, School of Biological Sciences, CH Waddington Building, The University of Edinburgh, United Kingdom

## Abstract

How cells choose between potential carbon sources is a classic example of cellular decision-making, and we know that many organisms prioritise glucose. Yet there has been little investigation of whether other sugars are also preferred, blinkering our view of carbon sensing. Here we study eukaryotic budding yeast and its growth on mixtures of palatinose, an isomer of sucrose, with other sugars. We find that yeast prioritise galactose over palatinose, but not sucrose or fructose, despite all three of these sugars being able to support faster growth than palatinose. Our results therefore disfavour carbon flux-sensing as the sole mechanism. By using genetic perturbations and transcriptomics, we show that repression is active and through Gal4, the master regulator of the GAL regulon. Cells enforce their preference for galactose over palatinose by preventing runaway positive feedback in the MAL regulon, whose genes enable palatinose catabolism. They do so both by repressing MAL11, the gene encoding the palatinose transporter, and by first expressing the isomaltases, IMA1 and IMA5, which cleave palatinose and so prevent its intracellular concentration becoming enough to induce further MAL expression. Our results demonstrate that budding yeast actively maintain a preference for carbon sources other than glucose and that such preferences have been selected by more than differences in growth rates. They imply that carbon-sensing strategies even in unicellular organisms are more complex than previously thought.

## Introduction

All cells respond to change. Understanding the strategies that they use to do so is fundamental because we expect these strategies to be more deeply conserved than how they are biochemically implemented [1, 2, 3], with different cell types realising the same strategy in different ways.

A classic example of decision-making is whether a cell consumes two available carbon sources either sequentially — often called diauxie [4] — or simultaneously. For both the bacterium *Escherichia coli* and the eukaryote *Saccharomyces cerevisiae*, glucose is preferred, and at sufficient concentrations, cells specialise their physiology to its consumption. For *S. cerevisiae*, cells both repress expression of genes for metabolising other carbon sources [5] and remove their transporters from the plasma membrane [6, 7, 8, 9]. Yet apart from glucose, budding yeast can consume at least six other sugars [10], and we know little about how or even whether cells discriminate between them.

We therefore do not have a clear picture of how yeast, one of the most studied eukaryotic cells, organise their carbon-sensing, a fundamental task that involves kinases conserved even in metazoans [11]. Although much regulation is known to impose the cells’ preference for glucose, it is unclear if similar complexity exists to enforce a hierarchy of preferences for all pairs of sugars or if control is more generic, perhaps through sensing of glycolytic flux as happens in *E. coli* [12, 13] or occurring passively through dilution because different sugars allow different growth rates [14].

Here we systematically investigate budding yeast’s decision-making on two sugars neither of which is glucose. Cells import sugars in two ways, via either hexose transporters or proton symporters [10]. If the same transporters import both sugars, the sugars may compete to bind the transporters [15]. We therefore chose pairs of sugars that require both types of import mechanisms, reasoning that such sugars are more likely to be independently regulated.

For the sugar requiring proton symport, we focused on palatinose, a disaccharide of glucose and fructose and a constituent of sugar cane and honey [16]. Palatinose is a substrate of the MAL regulon [16]. The laboratory strain BY4741, and its prototrophic antecedent FY4, both grow on palatinose but not on the more studied maltose [16], another disaccharide also imported by proton symporters. Palatinose is the only known substrate of these strains’ MAL regulons.

We found that budding yeast does have a sugar hierarchy beyond glucose, but it is complex. We observed diauxie in mixtures of galactose and palatinose, and too for glucose and palatinose, but not in mixtures of fructose or sucrose and palatinose. Combining genetic perturbations and transcriptomics, we show that cells implement their preference for galactose both by repressing the expression of MAL11, encoding the palatinose transporter, and by expressing the isomaltases, the enzymes that catabolise palatinose. Our results point not towards generic carbon-sensing, but towards specific regulation that actively enforces a sugar hierarchy.

## Results

### Cells growing in galactose-palatinose mixtures show diauxie

We used plate readers to characterise the cells’ growth, measuring the optical density (OD) and where appropriate the fluorescence of cultures. With the `omniplate` software package [17], we correct for the non-linear dependence of the OD on cell number [18] and for autofluorescence [19], use Gaussian processes to estimate growth rates over time [20], and automatically extract regions of exponential growth [21].

We observe diauxic-like growth for galactose-palatinose mixtures, similar to the expected diauxie [5, 22] that we also see in glucose-palatinose mixtures (Fig. 1C). Surprisingly, there is no obvious diauxie in fructose-palatinose or sucrose-palatinose mixtures, despite both allowing growth at a rate similar to that in glucose (Fig. 1B). We confirmed that the galactose-palatinose diauxie depends neither on the sugar concentrations (Fig. S1A–D) nor the pre-growth (Fig. S2A–C) and is not an artefact, with the cells consuming the ethanol or acetate generated by growing on galactose (Fig. S2D).

**Figure 1.**
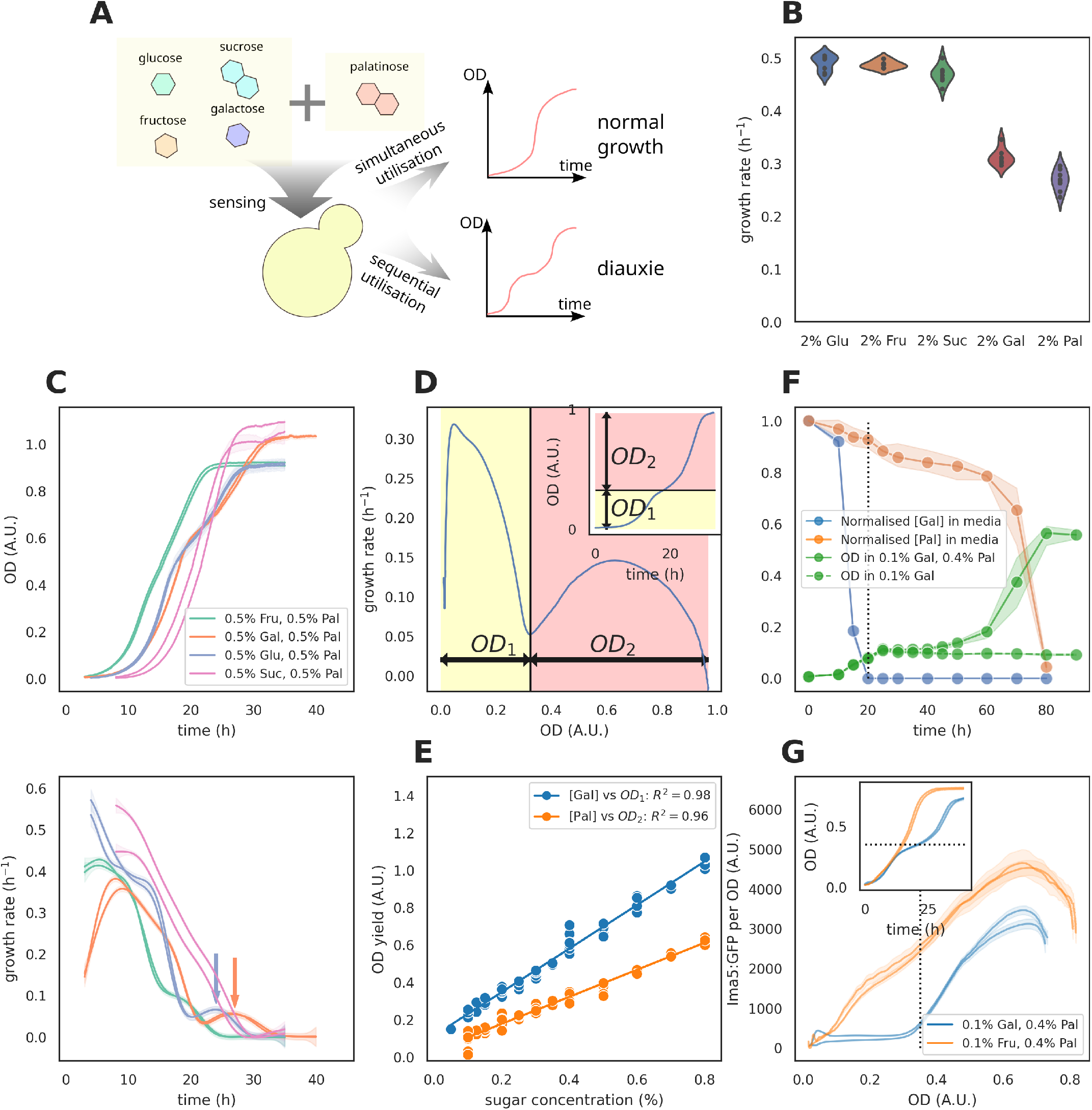
Cells consume galactose before palatinose. **(A)** We grow budding yeast cells in glucose-, fructose-, sucrose- and galactose-palatinose mixtures and observe the growth dynamics. **(B)**Budding yeast grows at different rates on different sugars, with palatinose supporting the slowest growth. **(C)** We observed diauxie in the growth dynamics of the wild-type prototrophic strain (FY4) in glucose- and galactose-palatinose mixtures. The arrows point to the second peak of growth rate in glucose- and galactose-palatinose mixtures. **(D)** To quantify the OD yield of each growth phase, we found the local minimum of the specific growth rate between the two maxima. If this minimum marks the end of growth phase 1 and the beginning of growth phase 2, then the OD yield of growth phase 1 (*OD*_1_) is the OD at the local minimum and the OD yield of growth phase 2 (*OD*_2_) is the difference between the final OD and *OD*_1_. **(E)** In galactose-palatinose mixtures, the OD yield of growth phase 1 linearly correlates with galactose concentrations, and the OD yield of growth phase 2 linearly correlates with palatinose concentrations. We find each data point using the method shown in (D). **(F)** Metabolomics data confirms that cells prioritise galactose over palatinose. We measure concentrations of extracellular galactose and palatinose by GC-MS, normalising by the values of the first time point (0 h). The OD of the samples is measured in a plate reader. Each data point represents the mean of three biological replicates and the shaded area their standard deviation. **(G)** The level of isomaltase Ima5:GFP per OD as a function of OD in fructose- and galactose-palatinose mixtures. Inset: the growth dynamics. The black dotted line marks the OD at which galactose is close to depletion. In Panels (C) and (G), each curve is from one biological replicate and the shaded area represents the standard deviation of two technical replicates.

Consistent with diauxie [23], the amount of growth in the two exponential periods of growth is proportional to the concentration of either galactose for the first phase or palatinose for the second phase. First we found the OD of the culture, OD_switch_, at the local minimum of the specific growth rate over time, which lies between the two maxima characteristic of diauxie (Fig. 1D). We then define the yield for the first growth period by the difference between OD_switch_ and OD_initial_ and the yield for the second period by the difference between OD_final_ and OD_switch_. The first yield linearly correlates with the galactose concentration and the second with the palatinose concentration (Fig. 1E).

Cells use two isomaltase enzymes, Ima1 and Ima5, to cleave palatinose [16]. Focusing on IMA5-GFP, we observed, as expected for diauxic growth, that Ima5 increases only after the first period of exponential growth in galactose-palatinose mixtures, but in contrast increases immediately in fructose-palatinose mixtures (Fig. 1G). We confirmed this behaviour at the single-cell level (Fig. S9).

Finally we grew cells in flasks and measured the extracellular sugar concentrations over time using metabolomics [24] (Fig. 1F). The galactose concentration vanished within 20 hours when approximately 90% of the palatinose was still present. The palatinose concentration, however, only quickly decreased during the second period of exponential growth.

Our results point towards a specific mechanism generating the galactose-palatinose diauxie. At similar concentrations, cells grow faster in sucrose and in fructose compared to palatinose (Fig. 1B), implying a higher glycolytic flux. Yet there is no apparent diauxie in both sucrose- and fructose-palatinose mixtures (Fig. 1C), inconsistent with either a general carbon flux-sensing mechanism [5, 25] or passive control through dilution [14].

### Active Gal4 delays the use of palatinose

To determine how intracellular galactose reduces the levels of Ima5 (Fig. 1H), we constitutively activated the GAL regulon by deleting the GAL80 gene. In the absence of galactose, Gal80 represses the activity of Gal4, the regulon’s master transcriptional regulator. Gal4 is always active in cells without Gal80 [26].

We observe that the *gal80*∆ strain either does not use or delays using palatinose in both galactose-palatinose and fructose-palatinose mixtures (Fig. 2A). This delay vanishes in a *gal80*∆ *gal4*∆ mutant and is absent in a *gal4*∆ (Fig. S3A): active Gal4 therefore likely prevents cells using palatinose.

**Figure 2.**
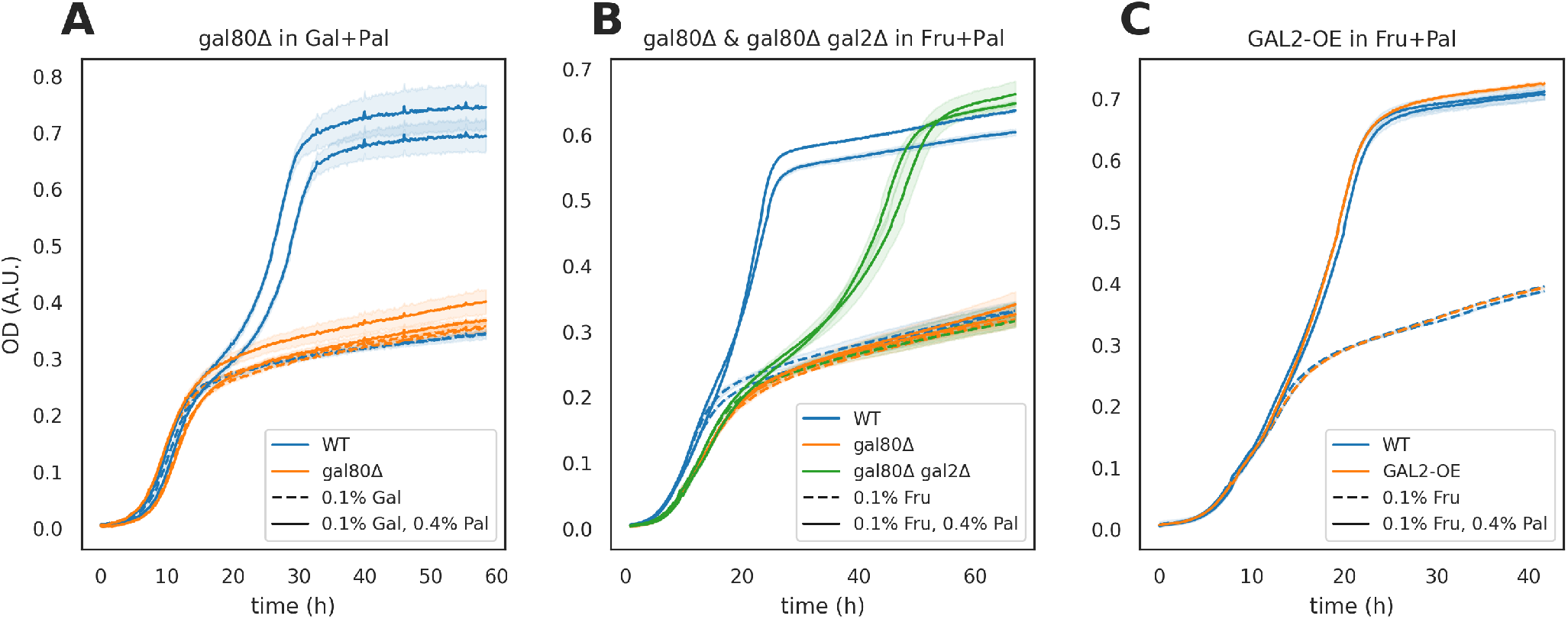
The Gal4 signal delays the use of palatinose. **(A)** Deleting GAL80 strongly delays growth in palatinose in galactose-palatinose mixture. **(B)** Deleting GAL80 strongly delays growth in palatinose in fructose-palatinose mixtures, and further deleting GAL2 partially alleviates the delay. **(C)** Over-expressing GAL2 with the CCW12 promoter (GAL2-OE) does generate a delay. In all panels, each curve represents one biological replicate and the shaded area represents the standard deviation of two technical replicates.

Gal4 induces the genes GAL1, GAL7, and GAL10, and this expression could deplete intracellular resources [27], preventing *gal80*∆ cells from expressing the MAL regulon in galactose-palatinose mixtures. Deleting the entire GAL1-10-7 locus in the *gal80*∆ mutant, however, did not change its phenotype (Fig. S3B).

Active Gal4 also induces expression of GAL2, which encodes galactose permease, a hexose transporter. Surprisingly, we found deleting GAL2 does allow the *gal80*∆ cells at least partially to consume palatinose (Fig. 2B), but over-expressing GAL2 in GAL80 cells fails to delay growth (Fig. 2C). Our results imply that active Gal4 and GAL2 together in some way impede cells from metabolising palatinose.

### Active Gal4 prevents MAL11 inducing

Gal4 is a transcriptional activator, and so we used RNA-seq to investigate how Gal4 in the *gal80*∆ mutant alters gene expression. We chose fructose as the other sugar: glucose is unsuitable because it represses GAL4 irrespective of Gal80’s presence [28] whereas fructose does not, and in fructose-palatinose mixtures, the wild-type strain co-consumes both fructose and palatinose in contrast to the *gal80*∆ mutant that consumes only fructose (Fig. 2B). We selected the concentration of fructose to make the growth of the wild-type and *gal80*∆ strains as similar as possible to reduce confounding transcriptional changes generated by differing growth rates [29].

The *gal80*∆ affects the expression of the two isomaltase genes and the palatinose transporter, MAL11 (Fig. 3A–C). With palatinose (lighter colours), the transcripts of the isomaltases in both the wild-type (blue) and the *gal80*∆ (orange) strains have increased by the mid-log time point, but while the wild-type’s keep increasing, those of the mutant stabilise. In contrast, the mutant’s MAL11 gene is never induced, unlike the wild-type’s.

**Figure 3.**
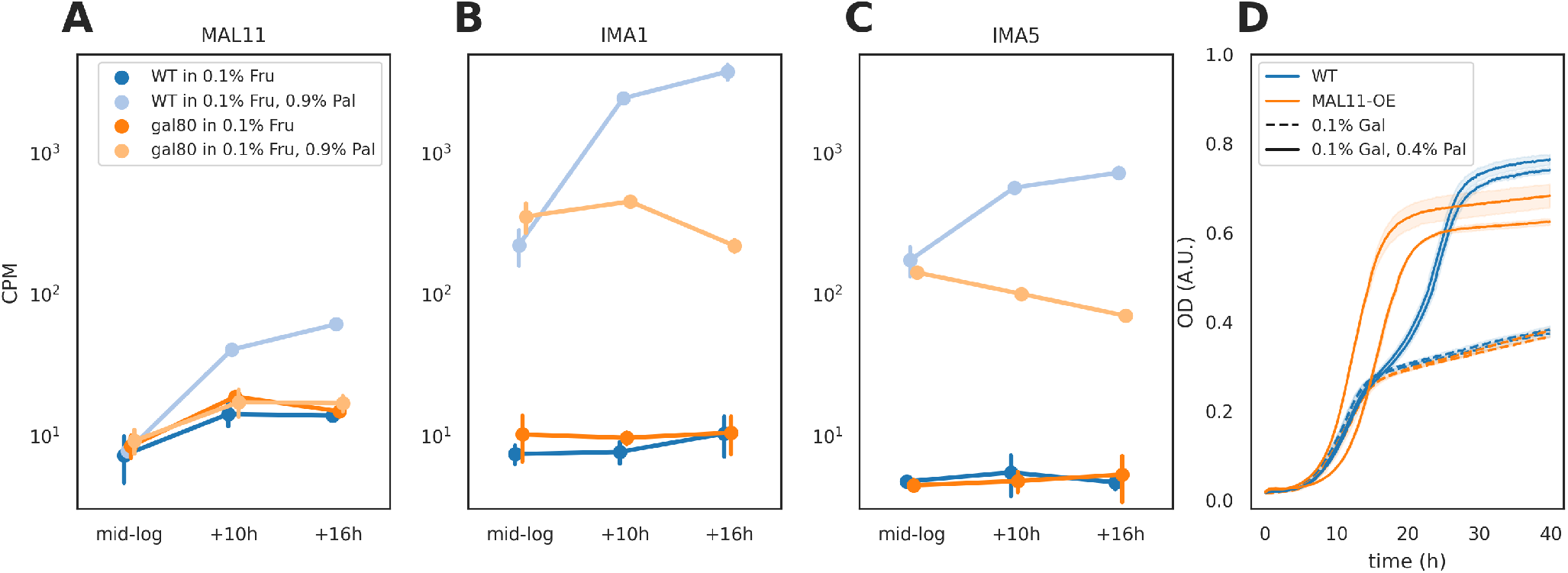
The Gal4 signal prevents the activation of MAL11 expression. **(A–C)** The count per million reads (CPM) of MAL11 (A), IMA1 (B) and IMA5 (C) transcripts. The error bar represents the standard deviation of three technical replicates. **(D)** Over-expressing MAL11 with the CCW12 promoter (MAL11-OE) in the wild-type abolishes the diauxie phenotype. Each curve represents one biological replicate. The shaded area represents the standard deviation of two technical replicates.

The MAL regulon has positive feedback: there are two transcriptional activators that induce expression of MAL11 in the presence of palatinose [16]. Higher levels of Mal11 generate more intracellular palatinose and so further activate MAL11 giving higher still levels of Mal11.

The RNA-seq results are consistent with active Gal4 repressing MAL11, either directly or indirectly, and so preventing runaway feedback in the MAL regulon. With the resulting low levels of Mal11, cells import enough palatinose to induce the isomaltase genes in the frucose-palatinose mixture, but not enough to generate positive feedback and induce MAL11’s expression. To test this hypothesis, we over-expressed MAL11 in both the wild-type and the *gal80*∆ strain and returned to galactose-palatinose mixtures. Consistently, both the diauxie in the wild type (Fig. 3D) and the deletion mutant’s delay vanish (Fig. S5A).

### The preference of galactose over palatinose results from the repressing GAL signal and early expression of the isomaltases

Although the MAL regulon has positive feedback and so can potentially exist in two states, one weakly and one strongly expressing, the isomaltases may prevent cells from reaching the strongly expressing state. If induced sufficiently quickly, the isomaltases may outcompete the regulon’s transcriptional activators for palatinose, cleaving it into fructose and glucose, and preventing runaway positive feedback. Consistently, for cells capable of metabolising maltose, over-expressing the maltase gene MAL12 generates a long lag when cells switch from glucose to maltose [30], likely because these high levels of Mal12 prevent cells inducing the MAL regulon. By inhibiting MAL11’s but not IMA1 and IMA5’s expression, Gal4 may use the same mechanism to prevent palatinose metabolism. Using mathematical modelling, we confirmed that cells can prioritise galactose over palatinose either by galactose strongly repressing the MAL regulon or by moderate repression with a negative feedback induced by higher levels of the isomaltases (SI & Fig. S6).

To test this role of the isomaltases, we decreased their levels by deleting an isomaltase gene. As expected, we find that *ima1*∆ cells lose diauxie in galactose-palatinose mixtures (Fig. 4A & B), although *ima5*∆ cells do not (Fig. S5C). This behaviour is still consistent however because deleting IMA1 likely decreases isomaltase concentrations more than deleting IMA5: in palatinose, IMA1’s transcript levels are five-fold higher than IMA5’s (Fig. 3B & C). Without IMA1, cells may have sufficiently low levels of isomaltase that the palatinose concentration necessary to generate runaway feedback becomes small enough that the feedback happens even with the low levels of Mal11 caused by galactose repression. Also in agreement, we find that a *gal80*∆ *ima1*∆ strain in galactose-palatinose mixtures loses the delay of the *gal80*∆ strain (Fig. S5B). Furthermore, deleting IMA1 decreases the lag and increases the growth rate in palatinose (Fig. 4C), consistent with runaway positive feedback in the MAL regulon enabling more palatinose import.

**Figure 4.**
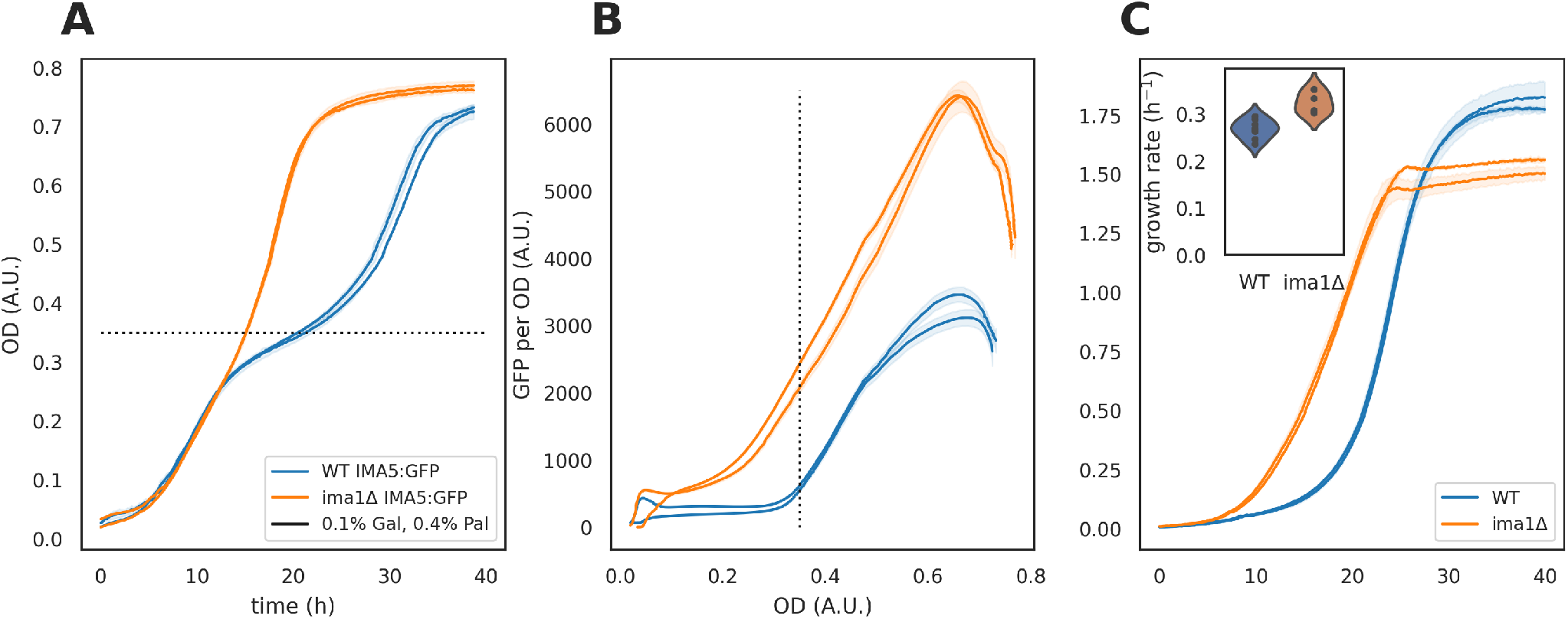
The preference of galactose over palatinose results from both repression by GAL and early expression of the isomaltases. **(A, B)** Deleting IMA1 from the wild-type strain abolishes diauxie in galactose-palatinose mixtures. The black dotted line marks the OD at which galactose is close to depletion. **(C)**Deleting IMA1 decreases the lag and increases the growth rate (inset) in 2% palatinose. In each panel, each curve represents one biological replicate and the shaded area represents the standard deviation of two technical replicates.

## Discussion

We have shown that budding yeast prioritises sugars other than glucose, consuming galactose before palatinose. Our results are consistent with early work suggesting cells prefer galactose over maltose [31]. Cells actively impose this preference, partly through the transcriptional regulator Gal4. In sucrose-palatinose and fructose-palatinose mixtures, however, we did not observe diauxie.

Our findings challenge current understanding. Although they are consistent with the observation that cells undergoing diauxie prioritise the carbon source allowing faster growth [25], they are inconsistent with its converse. Both fructose and sucrose enable faster growth than palatinose does, yet we observe no obvious diauxie in mixtures of palatinose with these sugars. Our results suggest further that cells prioritise carbon sources neither by a flux-sensing mechanism alone because faster growth typically implies a faster glycolytic flux [32] nor passively through dilution [14]. Cells likely combine a general flux-sensing mechanism, perhaps through AMP kinase and protein kinase A [33], with targeted regulation specific to carbon sources.

There is some evidence of this targeted regulation despite it being little studied. Both galactose [34] and fructose [35] repress the SUC2 gene, which encodes for the invertase enzyme used to metabolise sucrose and raffinose. Galactose also represses CYB2 [36], an oxidoreductase used to metabolise lactate.

We suspect that galactose prevents palatinose metabolism by stopping runaway positive feedback developing in the MAL regulon (Fig. 5). As cells consume galactose, they activate Gal4 and repress MAL11, the palatinose transporter. This repression together with early expression of the isomaltases, IMA1 and IMA5, prevent intracellular palatinose reaching sufficient concentrations to induce higher expression of MAL11. As cells exhaust galactose, however, Gal80 inactivates Gal4, and Gal4’s repression of MAL11 lifts, import of palatinose increases, and positive feedback develops.

**Figure 5.**
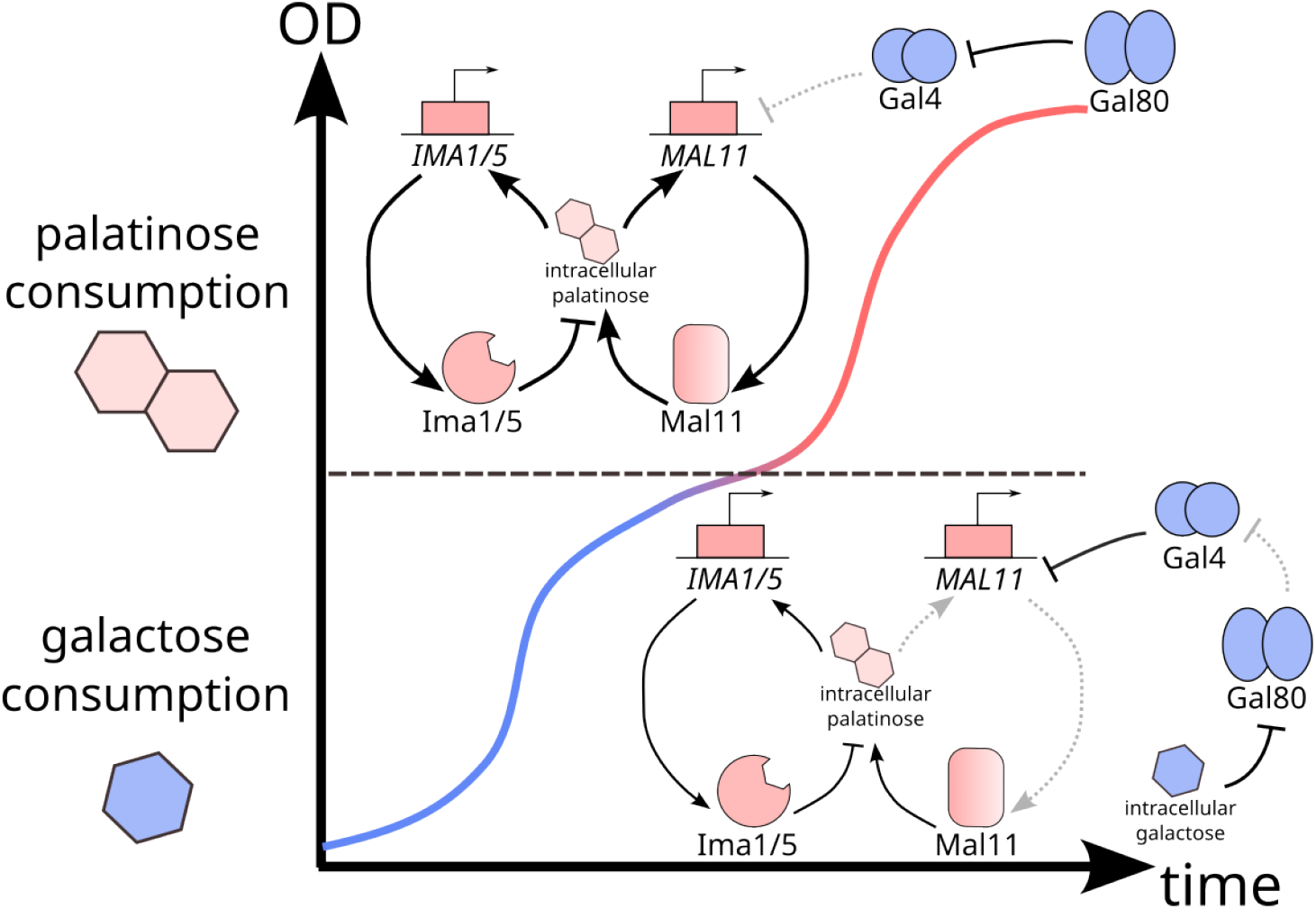
Cells use an active, specific mechanism to undergo galactose-palatinose diauxie. Initially they consume galactose. Active Gal4 represses MAL11, which together with the negative feedback through Ima1 and Ima5 prevents substantial positive feedback in the MAL regulon (greyed out arrows). When galactose runs out, Gal80 inactivates Gal4. The repression on MAL11 lifts, and the higher levels of the Mal11 transporters increase intracellular palatinose, further activating MAL11. Positive feedback in the MAL regulon becomes self-reinforcing, and cells consume palatinose.

Prioritising activation of the isomaltase genes may have been selected to prevent too much intracellular palatinose. Maltose, another substrate of the MAL regulon, is toxic at high intracellular concentrations and inhibits translation [37]. Its import, like palatinose’s, uses the proton-motive force and so may impose an energetic burden [38]. The regulon’s setup allows too flexibility in the decision-making: we showed that the loss of the IMA1 gene abolishes diauxie. IMA1, like most MAL genes, is near the telomeres, where gene loss and duplication are common [16].

We do not know how active Gal4 represses MAL11. Although Gal4 is reported to directly regulate only 12 genes [39, 40], our transcriptomic data imply that it affects the expression of a larger set, including the hexose transporters and genes controlling ribosome biogenesis (Fig. S7C & D), as well as the GAL regulon and other known non-GAL targets [39, 40, 41]. None of these genes, however, are transcription factors whose expression Gal4 could promote to repress MAL11.

A puzzling result is that deleting GAL2, the gene for galactose permease, partially alleviates the negative effects of deleting GAL80, allowing cells to re-consume palatinose in galactose-palatinose mixtures (Fig. 2C). Similarly, slow growth of the *gal80*∆ mutant in raffinose is also partly lifted by deleting GAL2 [42]. Perhaps removing GAL2 affects expression of nearby non-coding RNAs in the genome, such as the overlapping ncRNA SUT692 [43], whose function is unknown.

Our results suggest that budding yeast’s preference for glucose is not unique and that cells actively regulate to enforce preferences for other sugars, such as galactose. We do not understand why cells prioritise galactose and glucose over palatinose but not fructose or sucrose despite cells growing faster on fructose and sucrose than they do on galactose. This behaviour suggests that cells do more than maximise growth rates, even in laboratory conditions. Active regulation is presumably necessary because of intracellular constraints [44], but why these constraints should become alleviated in fructose and sucrose is unclear. Perhaps some of the behaviours we see are under only weak selection because yeast rarely encounter the corresponding combinations of sugars in the wild. More generic regulatory mechanisms may than suffice [45], such as control by SNF1 kinase, yeast’s equivalent of AMP kinase, and the repressor Mig1 [10]. Alternatively, for some sugars, competition may be fiercer than others, and so cells prioritise these sugars in an effort to starve competing organisms rather than for the sugars’ intrinsic values — such strategies can be evolutionarily stable [46].

Taken together, our findings imply that carbon-sensing is too important for cells to regulate with only generic mechanisms, and the onus now is both to delineate the decision-making strategies used and to determine how conserved they are across different species.

## Materials and Methods

### Strains and growth media

We list strains and constituents of the media used in Supplementary Tables 1 and 2. The BY4741-background strains are auxotrophic and the FY4-background strains are prototrophic [47]. Strains were pre-cultured in synthetic complete (SC) media supplemented with 2% (w/v) sodium pyruvate for two days before experiments, unless specified otherwise. We then diluted cultures six-fold six hours before an experiment with fresh SC media with 2% (w/v) sodium pyruvate to ensure cells are at exponential growth when the experiment begins. During an experiment, we grew auxotrophic strains in SC or LoFlo media and prototrophic strains in minimal media (Delft media) [48, 49], both supplemented with carbon sources.

### Creating yeast strains

We followed a standard protocol using lithium acetate and polyethylene glycol (PEG) to transform yeast [50]. Transformants were confirmed by colony PCR and Sanger sequencing (MRC Protein Phosphorylation and Ubiquintylation Unit, Dundee). We list all plasmids that we used in Supplementary Table 3. See also Supplementary Methods for multiplex CRISPR, which we used to delete the GAL1-10-7 locus.

### Growth assay in plate readers

We used plate readers (Tecan, Infinite M200 Pro or F200) to measure the dynamics of growth and fluorescence (Fig. 1A). Cells were grown in SC + 2% (w/v) sodium pyruvate in a 30 *◦*C shaking incubator at 180 rpm for about 40 hours and then diluted by six-fold 6–8 hours before the experiment. Before harvesting, we added 20 *µ*L 10x sugar stock or water to each well, and cultures of each strain were then centrifuged at 3500 rpm for 3 minutes and the supernatant removed. We washed cells using the appropriate media base once for experiments with SC or LoFlo and twice for experiments with Delft media. Cells were then re-suspended so that the initial OD was below 0.2 as measured by a spectrophotometer. Finally, we added 180 *µ*L re-suspended culture to each well to give a final volume of 200 *µ*L. We then moved the 96-well plate into the plate reader at 30 *◦*C with linear shaking at an amplitude of 6 mm and measurements taken every 10 minutes.

The plate-reader data are typically time series of 96 wells with both OD and fluorescence readings. We used a Python package, `omniplate` (version 0.9.92) [17], to analyse the data. Our typical pipeline is: (1) ignore any contaminated wells; (2) average over technical replicates and estimate the error; (3) subtract the OD and fluorescence background of the media; (4) correct the non-linearity between OD and the cell number when OD is high [18]; (5) estimate the specific growth rate (*d/dt* log *OD*) using a Gaussian process [20], along with other quantities such as maximal OD; (6) if fluorescence is measured, correct the auto-fluorescence using untagged cells and spectral unmixing [19]; (7) calculate the fluorescence reading per OD.

### Measuring sugar concentrations by Gas Chromatography – Mass Spectrometry (GC-MS)

#### Growing the cells and harvesting the spent media

We grew cells of the FY4 wild-type strain in SC+2% pyruvate in a 30 *◦*C shaking incubator at 180 rpm for about 40 hours and then diluted by six-fold six hours before the experiment. When the experiment began, we washed the cells twice with Delft media without carbon sources and then inoculated into 250 mL flasks with 25 mL Delft media supplemented with the desired concentrations of galactose and palatinose. The volume of inoculated cells was calculated to make the initial OD 0.05, and then we topped up the volume of each culture to 26 mL. The cultures were then incubated in a 30 *◦*C shaking incubator at 180 rpm.

To harvest the spent media, we sampled 1 mL of each culture into a 15 mL Falcon tube placed on ice and then immediately put the flasks back into the shaking incubator to minimise the impact of sampling. From each 1 mL sample, we transferred 2 *×*200 *µ*L samples into two wells of a 96-well microplate for OD measurement in a Tecan plate reader (Tecan, Infinite M200 Pro). The remaining volumes in the samples were centrifuged at 4000 rpm for 15 minutes at 4 *◦*C, and then we transferred 50 *µ*L of the supernatant into a GC vial and stored at -20 *◦*C. We harvested samples at 0, 10, 15, 20, 25, 30, 40, 50, 60, 70 and 80 h and measured the final OD at 90 h. In parallel, we measured with the same plate reader the OD of cultures in 0.1% galactose as a negative control.

#### Sample and standards preparation for sugar analysis

To the 50 *µ*L spent media, we added 5 *µ*L of the internal standard (3 mg/mL myristic acid d27 dissolved in water: methanol: isopropanol in a ratio of 2: 5: 2, v/v/v). The contents of the GC vial were evaporated to dryness in a Gene-Vac EZ-2 Elite evaporator, and trimethylsilylated with 50 *µ*L pyridine: N-methyl-N-trimethylsilyltrifluoroacetamide (1:4) for gas chromatography quadrupole time-of-flight mass spectrometry (GC/QTOF-MS) analysis of the sugars.

#### GC-MS analysis

The sugar concentrations were analysed on an Agilent 7890B gas chromatogram (GC) coupled to an Agilent 7200B quadrupole time-of-flight mass spectrometer (QTOF-MS) with GERSTEL multipurpose sampler (MPS) robotics (Anatune). Trimethysilylated samples (1 *µ*L) were injected at a split ratio of 10:1, with a split flow of 10 mL/min into a DB-5ms 40 m*×* 250 *µ*m *×* 0.25 *µ*m GC column (Agilent Technologies). We used helium as the carrier gas at a flow rate of 1 mL/min and set the inlet to 250 *◦*C and programmed the GC oven to 60 *◦*C for 1 min, followed by ramping at 10 *◦*C/min to 325 *◦*C, where it was held for 10 min. The ion source was set to 230 *◦*C, 35 *µ*A filament current, 70 eV electron energy, and we scanned the mass range of 60–900 m/z at an acquisition rate of 4 spectra/s with a solvent delay of 5 min. Total ion chromatograms and mass spectra were analysed using the Agilent MassHunter Qualitative Analysis B.10.00 software, and peak areas calculated using the Agile 2 integrator method.

### RNA measurements

#### Growing and harvesting cells

We harvested approximately four OD units of cells, by sampling *x* mL of each culture, such that the value of *OD · x* is around 4, and then centrifuging the cells at 3500 rpm for 3 minutes at 4 *◦*C. The supernatant was removed and the cell pellets stored in -80 *◦*C if RNA extraction did not immediately follow.

#### Extracting RNA

We adapted a column-based protocol in [51] to extract RNA. We thawed the cell pellets on ice and then resuspended with 400 *µ*L RNA binding buffer (Zymo, #R1013-2). The mixtures were then transferred to 2 mL screw cap tubes with zirconia beads inside, and then cell lysis performed using the PreCellys Evolution homogeniser (Bertin Instruments) — the samples were shaken at 6000 rpm for 10 seconds for three cycles, with a 10-second pause between each cycle, before being placed on ice for one minute. We repeated the shaking-ice bath process five further times. Then we centrifuged the lysates for 90 seconds and transferred each supernatant to a Zymo Spin IIICG column (Zymo, #C1006) and centrifuged again. We then mixed the flow through with 400 *µ*L 100% ethanol, transferred to a Zymo Spin IIC column (Zymo, #C1011), and centrifuged at 12000 *× g* for one minute. With the RNA being on the column, we discarded the flow through. We then sequentially added and centrifuged through the column 400 *µ*L DNA/RNA prep buffer (Zymo, #D7010-2), 600 *µ*L DNA/RNA wash buffer (Zymo, #D7010-3), and 400 *µ*L DNA/RNA wash buffer, discarding all flow through. Finally, we centrifuged the column again before adding 30 *µ*L nuclease free water (Ambion, #AM9937) to elute the RNA. All steps of centrifugation were performed at 12000 *× g* for one minute unless otherwise specified.

We measured the RNA concentrations with a spectrophotometer (DeNovix, #DS-11) and confirmed the quality of the RNA samples using a Fragment Analyzer (Advanced Analytical Technologies, Inc.) with the Standard Sensitivity RNA Analysis Kit (Agilent, #DNF-471).

### RNA-seq experiment

We grew cells of the wild-type FY4 and *gal80*∆ strains in SC+2% pyruvate in a 30 *◦*C shaking incubator at 180 rpm for about 40 hours and then diluted by six-fold six hours before the experiment began. Next the cells were washed twice with Delft media without carbon sources and then inoculated into 250 mL flasks with 25 mL Delft media supplemented with the desired concentrations of fructose and palatinose. We calculated the volume of inoculated cells to make an initial OD of 0.005 and topped up the volume of each culture to 26 mL. The cultures were incubated in a 30 *◦*C shaking incubator at 180 rpm.

We harvested samples at three time points: mid-log (at OD 0.3), 10 hours after mid-log, and 16 hours after mid-log (Fig. S4C).

Edinburgh Clinical Research Facility performed quality control, library preparation, and sequencing. They used a Fragment Analyser Automated Capillary Electrophoresis System (Agilent Technologies Inc, #5300) with the Standard Sensitivity RNA Analysis Kit (#DNF-471-0500) for quality control and an Qubit 2.0 Fluorometer (Thermo Fisher Scientific Inc, #Q32866) with the Qubit RNA broad range assay kit (#10210) for quantification. To quantify DNA contamination, an Qubit dsDNA HS assay kit (#Q32854) was used.

They generated libraries from 400 ng of each total RNA sample with the QuantSeq 3’ mRNA Library Prep Kit REV for Illumina (Lexogen Inc, #016) according to the manufacturer’s protocol. These libraries were then quantified by fluorometry with the Qubit dsDNA High Sensitivity assay and assessed for quality and fragment size with the Agilent Fragment Analyser with the SS NGS Fragment 1–6000 bp kit (#DNF-473-33).

They performed 2 *×* 50 bp paired-end sequencing on the NextSeq 2000 platform (Illumina Inc, #20038897) using NextSeq 1000/2000 P2 Reagents (100 cycles) v3 (#20046811), which produced 46.49 Gbp data. The data produced by the NextSeq 1000/2000 Control Software (Version 1.4.1.39716) was then automatically uploaded to BaseSpace (Illumina) and converted into FASTQ files.

We carried out RNA-seq alignment and quality control following Haynes *et al*. [52] using code written in Nextflow [53] (Fig. S4D) and available in a git repository: https://github.com/DimmestP/nextflow_paired_reads_pipeline. We list the software versions we used in Supplementary Table 4. We adapted the genome annotation file from the longest transcripts taken from Table S3 in [54], and for genes without an reported 3’UTR in [54], we assigned a default-length UTR of 125 nt as the median length is reported at 128 nt. We modified the annotations of some MAL genes — MAL32, IMA1, MAL11, and MAL12 — and some genes neighbouring a MAL gene — VTH1, HXT8, VTH2, and ALR2 — according to their actual 3’ ends from the reads in our experiment. We also added the annotation of ZNF1 (YFL052W), which was missing. The output of this pipeline is a 5697 *×* 36 table with raw counts, which we used for differential expression analysis with DESeq2 (version 1.34.0) [55]. We then defined the set of differentially expressed genes between two conditions by | log_2_ fold change| *>* 0.5 and the adjusted p-value *<* 0.05 for all three time points (Fig. S7A & Fig. S8). Both the adjusted p-value and the log_2_ fold change were calculated with DESeq2 [55].

## Data availability

The RNA-seq data are available on the Gene Expression Omnibus (GEO) of National Center of Biotechnology Information (NCBI) with accession number GSE240743.

## Acknowledgements

YH gratefully acknowledges financial support from the Darwin Trust and PSS from the BBSRC and the Leverhulme Trust (grant number RPG-2018–004). WD is supported by the Medical Research Council (grant number MR/N013166/1). EWJW is supported by a Sir Henry Dale Fellowship jointly funded by the Wellcome Trust and the Royal Society (208779/Z/17/Z). We thank Richard Clarke, Angie Fawkes, and Lee Murphy, for performing RNA-seq at the Genetics Core of the Edinburgh Wellcome Trust Clinical Research Facility, and Sam Haynes, for his help with the RNA-seq pipeline.

## References

[1] Perkins, T. J. and Swain, P. S. Strategies for cellular decision-making. Mol. Syst. Biol. 5(326), 1–15 (2009).

[2] Balázsi, G., Van Oudenaarden, A., and Collins, J. J. Cellular decision making and biological noise: from microbes to mammals. Cell 144(6), 910–925 (2011).

[3] Bizzarri, M., Brash, D. E., Briscoe, J., Grieneisen, V. A., Stern, C. D., and Levin, M. A call for a better understanding of causation in cell biology. Nat. Rev. Mol. Cell. Biol. 20(5), 261–262 (2019).

[4] Monod, J. Diauxie et respiration au cours de la croissance des cultures de B. coli. Ann. Inst. Pasteur 68, 548–550 (1942).

[5] Gancedo, J. M. Yeast Carbon Catabolite Repression. Microbiol. Mol. Biol. Rev. 62(2), 334–361 (1998).

[6] Medintz, I., Jiang, H., Han, E. K., Cui, W., and Michels, C. A. Characterization of the glucose-induced inactivation of maltose permease in Saccharomyces cerevisiae. J. Bacteriol. 178(8), 2245–2254 (1996).

[7] Paiva, S., Vieira, N., Nondier, I., Haguenauer-Tsapis, R., Casal, M., and Urban-Grimal, D. Glucose-induced Ubiquitylation and Endocytosis of the Yeast Jen1 Transporter. J. Biol. Chem. 284(29), 19228–19236, July (2009).

[8] Horak, J. and Wolf, D. H. The ubiquitin ligase SCFGrr1 is required for Gal2p degradation in the yeast Saccharomyces cerevisiae. Biochem. Biophys. Res. Commun. 335(4), 1185–1190, October (2005).

[9] Hatanaka, H., Omura, F., Kodama, Y., and Ashikari, T. Gly-46 and His-50 of Yeast Maltose Transporter Mal21p Are Essential for Its Resistance against Glucose-induced Degradation. J. Biol. Chem. 284(23), 15448–15457, June (2009).

[10] Horák, J. Regulations of sugar transporters: Insights from yeast. Curr. Genet. 59(1-2), 1–31 (2013).

[11] Chantranupong, L., Wolfson, R. L., and Sabatini, D. M. Nutrient-sensing mechanisms across evolution. Cell 161(1), 67–83 (2015).

[12] Aidelberg, G., Towbin, B. D., Rothschild, D., Dekel, E., Bren, A., and Alon, U. Hierarchy of non-glucose sugars in Escherichia coli. BMC Syst. Biol. 8, 133–133 (2014).

[13] Okano, H., Hermsen, R., Kochanowski, K., and Hwa, T. Regulation underlying hierarchical and simultaneous utilization of carbon substrates by flux sensors in Escherichia coli. Nat. Microbiol. 5(1), 206–215 (2020).

[14] Narang, A. and Pilyugin, S. S. Bacterial gene regulation in diauxic and non-diauxic growth. J. Theor. Biol. 244(2), 326–348 (2007).

[15] Escalante-Chong, R., Savir, Y., Carroll, S. M., Ingraham, J. B., Wang, J., Marx, C. J., and Springer, M. Galactose metabolic genes in yeast respond to a ratio of galactose and glucose. Proc. Natl. Acad. Sci. U.S.A. 112(5), 1636–1641 (2015).

[16] Brown, C. A., Murray, A. W., and Verstrepen, K. J. Rapid Expansion and Functional Divergence of Subtelomeric Gene Families in Yeasts. Curr. Biol. 20(10), 895–903 (2010).

[17] Montaño-Gutierrez, L. F., Moreno, N. M., Farquhar, I. L., Huo, Y., Bandiera, L., and Swain, P. S. Analysing and meta-analysing time-series data of microbial growth and gene expression from plate readers. PLoS Comput. Biol. 18(5), e1010138, May (2022).

[18] Stevenson, K., McVey, A. F., Clark, I. B., Swain, P. S., and Pilizota, T. General calibration of microbial growth in microplate readers. Sci. Rep. 6(November), 4–10 (2016).

[19] Lichten, C. A., White, R., Clark, I. B., and Swain, P. S. Unmixing of fluorescence spectra to resolve quantitative time-series measurements of gene expression in plate readers. BMC Biotechnol. 14 (2014).

[20] Swain, P. S., Stevenson, K., Leary, A., Montano-Gutierrez, L. F., Clark, I. B., Vogel, J., and Pilizota, T. Inferring time derivatives including cell growth rates using Gaussian processes. Nat. Commun. 7(May), 1–8 (2016).

[21] Huo, Y., Li, H., Wang, X., Du, X., and Swain, P. S. Nunchaku: Optimally partitioning data into piece-wise linear segments. bioRxiv, 2023–05 (2023).

[22] New, A. M., Cerulus, B., Govers, S. K., Perez-Samper, G., Zhu, B., Boogmans, S., Xavier, J. B., and Verstrepen, K. J. Different Levels of Catabolite Repression Optimize Growth in Stable and Variable Environments. PLoS Biol. 12(1), 17–20 (2014).

[23] Monod, J. THE PHENOMENON OF ENZYMATIC ADAPTATION And Its Bearings on Problems of Genetics and Cellular Differentiation. Growth Symp. XI(12), 223–289 (1947).

[24] Moses, T., Thevelein, J. M., Goossens, A., and Pollier, J. Comparative analysis of CYP93E proteins for improved microbial synthesis of plant triterpenoids. Phytochemistry 108, 47–56, December (2014).

[25] Okano, H., Hermsen, R., and Hwa, T. Hierarchical and simultaneous utilization of carbon substrates: Mechanistic insights, physiological roles, and ecological consequences. Curr. Opin. Microbiol. 63, 172–178, October (2021).

[26] Lohr, D., Venkov, P., and Zlatanova, J. Transcriptional regulation in the yeast GAL gene family: A complex genetic network. FASEB J. 9(9), 777–787 (1995).

[27] Malakar, P. and Venkatesh, K. V. GAL regulon of Saccharomyces cerevisiae performs optimally to maximize growth on galactose. FEMS Yeast Res. 14(2), 346–356 (2014).

[28] Ricci-Tam, C., Ben-Zion, I., Wang, J., Palme, J., Li, A., Savir, Y., and Springer, M. Decoupling transcription factor expression and activity enables dimmer switch gene regulation. Science 372(6539), 292–295, April (2021).

[29] Brauer, M. J., Huttenhower, C., Airoldi, E. M., Rosenstein, R., Matese, J. C., Gresham, D., Boer, V. M., Troyanskaya, O. G., and Botstein, D. Coordination of Growth Rate, Cell Cycle, Stress Response, and Metabolic Activity in Yeast. Mol. Biol. Cell 19(1), 352–367, January (2008).

[30] Cerulus, B., Jariani, A., Perez-Samper, G., Vermeersch, L., Pietsch, J., Crane, M. M., New, A. M., Gallone, B., Roncoroni, M., Dzialo, M. C., Govers, S. K., Hendrickx, J., Galle, E., Coomans, M., Berden, P., Swain, P. S., and Verstrepen, K. J. Transition between fermentation and respiration determines history-dependent behavior in fluctuating carbon sources. eLife 7 (2018).

[31] Spiegelman, S. and Dunn, R. INTERACTIONS BETWEEN ENZYME-FORMING SYSTEMS DURING ADAPTATION. J. Gen. Physiol. 31(2), 153–173, November (1947).

[32] Huberts, D. H. E. W., Niebel, B., and Heinemann, M. A flux-sensing mechanism could regulate the switch between respiration and fermentation. FEMS Yeast Res. 12(i), 118–128 (2012).

[33] Broach, J. R. Nutritional control of growth and development in yeast. Genetics 192(1), 73–105 (2012).

[34] Gancedo, J. M., Flores, C.-L., and Gancedo, C. The repressor Rgt1 and the cAMP-dependent protein kinases control the expression of the SUC2 gene in Saccharomyces cerevisiae. Biochim. Biophys. Acta 1850(7), 1362–1367, July (2015).

[35] De Winde, J. H., Crauwels, M., Hohmann, S., Thevelein, J. M., and Winderickx, J. Differential Requirement of the Yeast Sugar Kinases for Sugar Sensing in Establishing the Catabolite-Repressed State. Euro. J. Biochem. 241(2), 633–643 (1996).

[36] Lodi, T., Donnini, C., and Ferrero, I. Catabolite repression by galactose in overexpressed GAL4 strains of Saccharomyces cerevisiae. Microbiology 137(5), 1039–1044 (1991).

[37] Hatanaka, H., Mitsunaga, H., and Fukusaki, E. Inhibition of Saccharomyces cerevisiae growth by simultaneous uptake of glucose and maltose. J. Biosci. Bioeng. 125(1), 52–58, January (2018).

[38] Eames, M. and Kortemme, T. Cost-Benefit Tradeoffs in Engineered lac Operons. Science 339(August 2011), 911–915 (2012).

[39] Ren, B., Robert, F., Wyrick, J. J., Aparicio, O., Jennings, E. G., Simon, I., Zeitlinger, J., Schreiber, J., Hannett, N., Kanin, E., Volkert, T. L., Wilson, C. J., Bell, S. P., and Young, R. A. Genome-Wide Location and Function of DNA Binding Proteins. Science 290(5500), 2306–2309, December (2000).

[40] Rhee, H. S. and Pugh, B. F. Comprehensive Genome-wide Protein-DNA Interactions Detected at Single-Nucleotide Resolution. Cell 147(6), 1408–1419, December (2011).

[41] Zheng, W., Xu, H. E., and Johnston, S. A. The cysteine-peptidase bleomycin hydrolase is a member of the galactose regulon in yeast. J. Biol. Chem. 272(48), 30350–30355, November (1997).

[42] Ideker, T., Thorsson, V., Ranish, J. A., Christmas, R., Buhler, J., Eng, J. K., Bumgarner, R., Goodlett, D. R., Aebersold, R., and Hood, L. Integrated Genomic and Proteomic Analyses of a Systematically Perturbed Metabolic Network. Science 292(5518), 929–934, May (2001).

[43] Parker, S., Fraczek, M., Wu, J., Shamsah, S., Manousaki, A., Dungrattanalert, K., Almeida, R., Estrada-Rivadeneyra, D., Omara, W., Delneri, D., and O’Keefe, R. A resource for functional profiling of noncoding RNA in the yeast Saccharomyces cerevisiae. RNA 23, May (2017).

[44] Bruggeman, F. J., Planqué, R., Molenaar, D., and Teusink, B. Searching for principles of microbial physiology. FEMS Microbiol. Rev. 44(6), 821–844, November (2020).

[45] Ammar, E. M., Wang, X., and Rao, C. V. Regulation of metabolism in Escherichia coli during growth on mixtures of the non-glucose sugars: arabinose, lactose, and xylose. Sci. Rep. 8(1), 609 (2018).

[46] Josephides, C. and Swain, P. S. Predicting metabolic adaptation from networks of mutational paths. Nat. Commun. 8(1), 685 (2017).

[47] Baker Brachmann, C., Davies, A., Cost, G. J., Caputo, E., Li, J., Hieter, P., and Boeke, J. D. Designer deletion strains derived from Saccharomyces cerevisiae S288C: A useful set of strains and plasmids for PCR-mediated gene disruption and other applications. Yeast 14(2), 115–132 (1998).

[48] Verduyn, C., Postma, E., Scheffers, W. A., and van Dijken, J. P. . Physiology of Saccharomyces Cerevisiae in Anaerobic Glucose-Limited Chemostat Cultures. Microbiology 136(3), 395–403 (1990).

[49] Verduyn, C., Postma, E., Scheffers, W. A., and Van Dijken, J. P. Effect of benzoic acid on metabolic fluxes in yeasts: A continuous-culture study on the regulation of respiration and alcoholic fermentation. Yeast 8(7), 501–517 (1992).

[50] Gietz, R. D. and Woods, R. A. Transformation of yeast by lithium acetate/single-stranded carrier DNA/polyethylene glycol method. Meth. Enzymol. 350, 87–96 (2002).

[51] Auxillos, J., Bayne, R., and Wallace, E. RNA extraction with spin columns from yeast cells grown on 12-column deep well plates. protocols.io, dx.doi.org/10.17504/protocols.io.beetjben, August (2021).

[52] Haynes, S., Auxillos, J., Danecka, W., Jain, A., Alibert, C., and Wallace, E. Limitations of composability of cis-regulatory elements in messenger RNA. bioRxiv, 2021.08.12.455418, April (2022).

[53] Di Tommaso, P., Chatzou, M., Floden, E. W., Barja, P. P., Palumbo, E., and Notredame, C. Nextflow enables reproducible computational workflows. Nat. Biotechnol. 35(4), 316–319, April (2017).

[54] Pelechano, V., Wei, W., and Steinmetz, L. M. Extensive transcriptional heterogeneity revealed by isoform profiling. Nature 497(7447), 127–131, May (2013).

[55] Love, M. I., Huber, W., and Anders, S. Moderated estimation of fold change and dispersion for RNA-seq data with DESeq2. Genome Biol. 15(12), 550, December (2014).

